# Highly Pathogenic Avian Influenza A (H5N1) Clade 2.3.2.1a virus infection in domestic cats, India, 2025

**DOI:** 10.1101/2025.02.23.638954

**Authors:** Ashwin Ashok Raut, Ashutosh Aasdev, Naveen Kumar, Anubha Pathak, Adhiraj Mishra, Prakriti Sehgal, Atul K Pateriya, Megha Pandey, Sandeep Bhatia, Anamika Mishra

## Abstract

In January 2025, the highly pathogenic avian influenza A(H5N1) virus clade 2.3.2.1a infection was detected in domestic cats and whole-genome sequencing of two cat H5N1 isolates was performed using the Oxford Nanopore MinION sequencing platform. Phylogenetic analysis revealed the circulation of triple reassortant viruses in cats. Although cat viruses lacked classic mammalian adaptation markers they carried mutations associated with enhanced polymerase activity in mammalian cells and increased affinity for α2–6 sialic acid receptor suggesting their potential role in facilitating infection in cats. The identification of reassortant HPAI H5N1 clade 2.3.2.1a viruses in domestic cats in India highlights the urgent need for enhanced surveillance in domestic poultry, wild birds, and mammals, including humans, to track genomic diversity and molecular evolution of circulating strains.

## Main text

Highly pathogenic avian influenza (HPAI) H5Nx viruses continue to pose a significant threat to wild birds, poultry, and mammals globally. Of them, HPAI H5N1 clade 2.3.4.4b viruses are of particularly alarming because of their capability of spillover into several mammals, including cats and their high genetic elasticity for reassortment (Plaza et al., 2024). While sporadic infection of cats with HPAI H5N1 has previously been reported, since 2020, clade 2.3.4.4b has been documented in large number of cats exhibiting severe neurologic and respiratory symptoms across at least five countries; France (Briand et al., 2023), Italy (Moreno et al., 2023), Poland (Rabalski et al., 2023), South Korea (Lee et al., 2024), and USA (Burrough et al., 2024). While clade 2.3.4.4b viruses have been the dominant circulating strains worldwide since 2020, several outbreaks in poultry caused by clade 2.3.4.4b and clade 2.3.2.1a viruses continues to be reported in India, raising concerns about potential spillover to mammals (WOAH, 2025). However, to date, only two human infections due to H5N1 clade 2.3.2.1a have been documented in India (Potdar et al., 2022; Deng et al., 2024), and no cases of HPAI H5N1 infection in cats have been reported. Here, we present the first documented cases of HPAI H5N1 clade 2.3.2.1a virus infection in domestic cats in India.

On 16^th^ and 24^th^ Jan, 2025, Indian Council Agricultural Research-National Institute of High Security Animal Diseases, Bhopal, India, received samples (whole blood, serum, and nasal swabs) of 4 and 3 cats respectively for virological testing (Appendix Table 1). These samples were submitted by the Veterinary Hospital Chhindwara, Madhya Pradesh, India. The cats, housed in separate houses within the same locality, exhibited clinical signs including pyrexia (102-104°F), anorexia and lethargy. Specific real-time reverse transcription PCR (RT-qPCR) targeting the matrix and H5 genes of Influenza A virus confirmed the presence of the H5Nx virus genome in samples from three cats (Appendix). All H5Nx positive cats died within 1-3 days after sampling. The viruses were successfully isolated from the blood samples of these cats using specific pathogen-free embryonated chicken eggs and subsequently, whole-genome sequencing of two viruses was performed from the allantoic fluid using the Oxford Nanopore MinION sequencing platform (Appendix).

The complete genome sequences are deposited in the NCBI GenBank database (accession nos. PV162485-PV162500). Analysis of hemagglutinin (HA) gene revealed the presence of a multiple basic amino acid cleavage site (PQKERRRKR*GLF) that, confirmed their classification as HPAI viruses. Genome analysis confirmed the viruses to be H5N1 subtype and with a single mutation in the PB1 gene between the two cat isolates: A/Cat/Chhindwara/ZD25-1/2025 (D383) and A/Cat/Chhindwara/ZD25-2/2025 (E383).

To identify the closest matching sequences in the entire GISAID database (https://platform.epicov.org/epi3/frontend#16d9ee), we conducted a BLAST analysis on all eight gene segments of the cat HPAI H5N1 viruses. The results showed a close match to an HPAI H5N1 detected in a traveler returning to Australia from India in 2024 (A/Victoria/149/2024, GISAID accession no. EPI_ISL_19156871; https://www.gisaid.org). All gene segments closely matched this strain, except for the M gene (Appendix Table 2). A whole-genome comparison between these cat viruses and A/Victoria/149/2024 showed 99.2% homology, with 27 mutations distributed across different gene segments, including a PB1-F2 (66S) marker associated with enhanced replication and virulence of H5N1 viruses (Schmolke et al., 2011) (Appendix Table 3) indicating their distinctiveness.

To further investigate the genetic relationships of the cat HPAI H5N1 sequences, we retrieved highly similar sequences and representative sequences from different clades (2.3.2.1, 2.3.2.1a, 2.3.2.1c, 2.3.4.4b) from the GISAID database (Elbe and Buckland-Merrett, 2017). These sequences were aligned using MAFFT v.7.475 (Katoh and Standley, 2013), and a phylogenetic tree was constructed using the TIM+F+G4 model, based on the Bayesian Information Criterion, in IQ-TREE v. 2 (Minh et al., 2020). Phylogenetic analysis indicated that A/Cat/Chhindwara/ZD25-1/2025(H5N1) and A/Cat/Chhindwara/ZD25-2/2025(H5N1) are reassortant viruses. Four gene segments (HA, NA, NP, and NS) were closely related to HPAI H5N1 clade 2.3.2.1a viruses circulating in Bangladesh, while the remaining four segments (PB2, PB1, PA, and MP) clustered with clade 2.3.4.4b viruses (Figure 1 and Appendix Figures 1-4). Notably, the matrix (MP) segment clustered with an HPAI H5N1 clade 2.3.4.4b virus detected in a wild bird in South Korea (A/Bean goose/Korea/21WC198/2022), while the polymerase gene complex (PB2, PB1, and PA) grouped with low pathogenicity avian influenza (LPAI) H5N1 clade 2.3.4.4b viruses that have been detected in poultry and wild birds in Asia since 2022 (Appendix Figures 1-4). These findings suggest that both HPAI and LPAI strains may have acted as intermediaries in donating internal genes to the HPAI H5N1 clade 2.3.2.1a viruses detected in this study. It may be noted that H5N1 has been recently detected in wild felines (WOAH, 2025) and in poultry of the adjoining region in India (WOAH, 2025). However, a comprehensive understanding of the spatiotemporal epidemiology across avian and mammalian hosts remains limited due to the paucity of H5N1 complete genomes from India. Currently, only three full genomes of H5N1 clade 2.3.4.4b and 27 of H5N1 clade 2.3.2.1a viruses are available in GISAID, highlighting the need for expanded surveillance and genomic characterization

**Figure 1:**
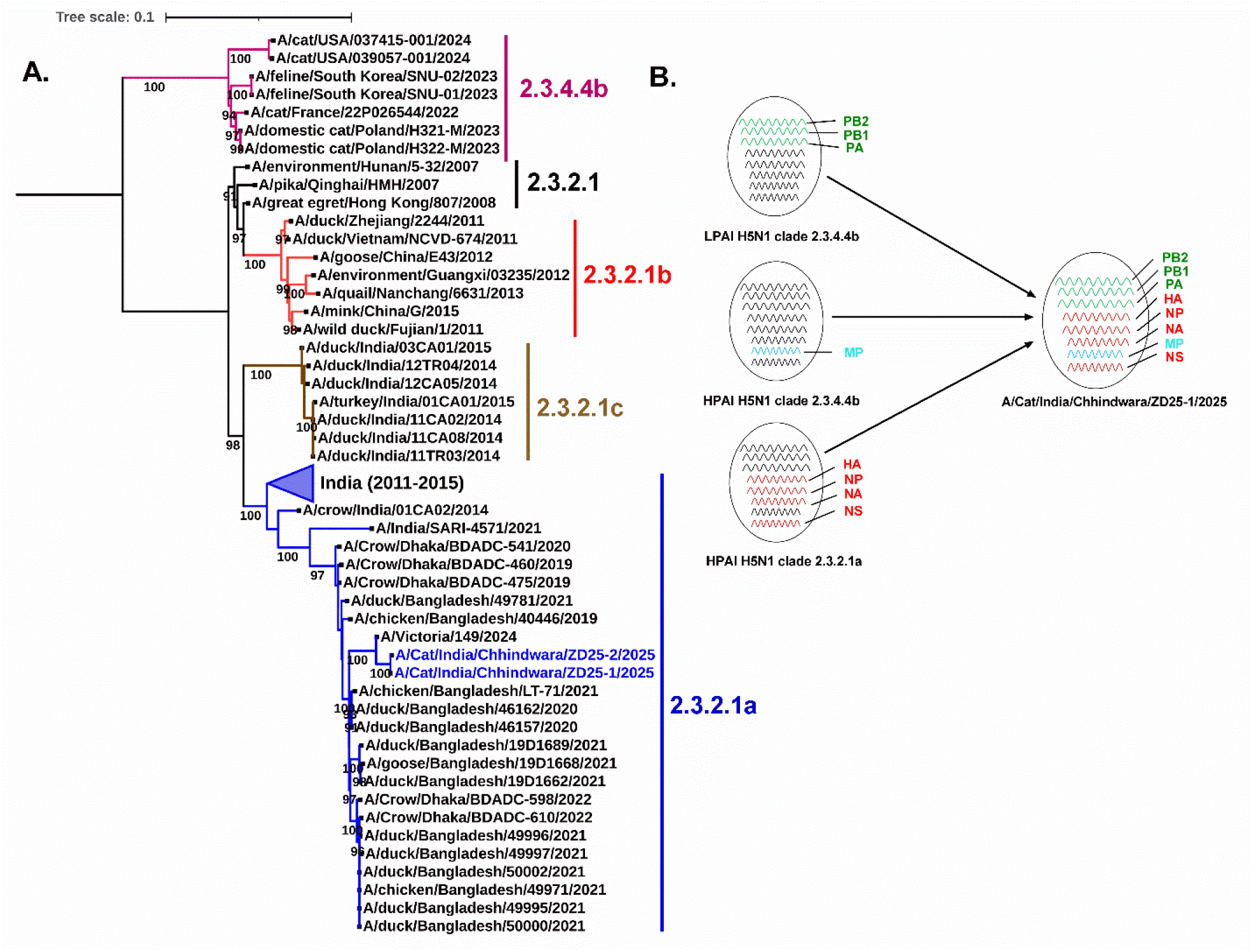
Evolutionary origins of cat HPAI H5N1 viruses detected in India. (A) Maximum-likelihood trees of the haemagglutinin (HA) gene for two cat HPAI H5 isolates — A/Cat/Chhindwara/ZD25-1/2025 and A/Cat/Chhindwara/ZD25-2/2025— demonstrating their phylogenetic relationships with BLAST sampled sequences and representative clade viruses (2.3.2.1, 2.3.2.1a, 2.3.2.1b, 2.3.2.1c, and 2.3.4.4b). The scale bar represents the number of nucleotide substitutions per site for HA gene. Bootstrap values exceeding 90% are shown. (B) Reassortant nature of the cat isolate-A/Cat/Chhindwara/ZD25-1/2025, inferred through a comprehensive phylogenetic analysis of all gene segments (Appendix Figures 1-4). HPAI, high pathogenicity avian influenza; LPAI, low pathogenicity avian influenza; PB2, polymerase basic 2; PB1, polymerase basic 1; PA, polymerase acidic; HA, hemagglutinin; NP, nucleoprotein; MP, matrix protein; NS, nonstructural protein.

Furthermore, we employed FluServer (https://platform.epicov.org/epi3/frontend#2fe8eb) to assess the phenotypic characteristics of all gene segments of the cat HPAI H5N1 viruses, focusing on mammalian adaptation, polymerase activity, virulence, and antiviral susceptibility. The analysis showed that the viruses exhibit a preference for avian α2–3 like sialic acid receptor binding, with key residues E190, G225, Q226, and G228. They remained susceptible to antiviral drugs, including oseltamivir, zanamivir, peramivir, and laninamivir. Although the viruses lacked classic mammalian adaptation markers in PB2 (E627K and D701N), they carried mutations associated with enhanced polymerase activity in mammalian cells (PB2-I292V, K389R, M676A; PA-K615R), and increased affinity for α2–6 sialic acid receptor (HA5 numbering-V210I), suggesting their potential role in facilitating infection in cats.

The identification of reassortant HPAI H5N1 clade 2.3.2.1a viruses in domestic cats in India highlights the urgent need for enhanced surveillance in domestic poultry, wild birds, and mammals, including pets and humans, to unravel the underlying genomic diversity and molecular evolution of H5N1 in India.

## Declaration

The views, findings, interpretations, or conclusions in the research paper do not necessarily represent the views of the institution, funding body, or any other affiliated organization.

## Acknowledgements

We are thankful to the Indian Council of Agricultural Research, New Delhi, and the Director, Indian Council of Agricultural Research–National Institute of High Security Animal Diseases, Bhopal, for providing necessary facilities to carry out this work. We are thankful to the Department of Animal Husbandry and Dairying Govt of Madhya Pradesh India for providing the clinical samples used as part of this study. We gratefully acknowledge the authors and the originating and submitting laboratories for the sequences from the Global Initiative on Sharing Avian Influenza Data EpiFlu database.

We acknowledge funding by Indian Council of Agricultural Research, New Delhi and the Department of Animal Husbandry, Dairying and Fisheries, Ministry of Agriculture and Farmers Welfare, Government of India under Central Disease Diagnostic Laboratory Grant

