## Appendix for "Highly Pathogenic Avian Influenza A (H5N1) Clade 2.3.2.1a virus infection in domestic cats, India, 2025"

### Supplementary Methods

#### Cats H5N1 whole genome sequencing on the Oxford Nanopore MinION platform

The whole genome sequencing of two cats H5N1 viruses were generated from the allantoic fluid on the Oxford Nanopore MinION sequencing platform. Briefly, a multisegment RT-PCR amplification was performed to amplify the complete genome segments of the H5N1 viruses using the OPTI primer set (1) and SuperScript III One-Step RT-PCR System with Platinum Taq DNA Polymerase, Invitrogen, USA. The resulting PCR products were purified, quantified, and 100 ng from each sample was utilized to construct a barcoded sequencing library with the PCR Barcoding Kit (SQK-PBK004). Sequencing was conducted on an Oxford Nanopore Technologies MinION device using an R9.4.1 flow cell, generating approximately 1.74 Gb data. Dorado v0.7.3 was used to basecall and demultiplex the data. Porechop ABI v0.5.0 (2) was used to trim the adapters and primer sequences. The genome segments were assembled using Iterative Refinement Meta-Assembler (IRMA), v1.2.0 (3). The reads were mapped to the consensus sequences using minimap2 v2.28 (4) to manually verify the sequences.

**Appendix Table 1. Cat clinical samples details received from the Veterinary Hospital Chhindwara, Madhya Pradesh, India.**

| Date of sample receipt | Place of collection | Animal Details | Sample Details | Lab Accession No | RT-qPCR test result | |
|---|---|---|---|---|---|---|
| | | | | | IAV Matrix | H5 subtype |
| January 16, 2025 | Chhindwara, Madhya Pradesh | Cat (Persian, 2.5 yr, female) | Blood | ZD/25/1-Bd | **Positive** | **Positive** |
| | | | Nasal Swab | ZD/25/1-NS | **Positive** | Negative |
| | | Cat (Persian, 14 mth, male) | Blood | ZD/25/2-Bd | **Positive** | **Positive** |
| | | | Nasal Swab | ZD/25/2-NS | Negative | H5 subtyping was not done for samples negative for Influenza A matrix |
| | | Cat (Non-Descript, 5 mth, female) | Blood | ZD/25/3-Bd | Negative | |
| | | | Nasal Swab | ZD/25/3-NS | Negative | |
| | | Cat (Non-Descript, 1yr, female) | Blood | ZD/25/4-Bd | Negative | |
| | | | Nasal Swab | ZD/25/4-NS | Negative | |
| January 24, 2025 | Chhindwara, Madhya Pradesh | Cat (Non-Descript, 6 mth, male) | Blood | ZD/25/5-Bd | **Positive** | Negative |
| | | | Serum | ZD/25/5-Se | **Positive** | Negative |
| | | | Nasal Swab | ZD/25/5-NS | Negative | Negative |
| | | Cat (Non-Descript, 1 yr, female) | Blood | ZD/25/6-Bd | Negative | Negative |
| | | | Serum | ZD/25/6-Se | Negative | Negative |
| | | | Nasal Swab | ZD/25/6-NS | Negative | Negative |
| | | Cat (Non-Descript, 1 yr, male) | Blood | ZD/25/7-Bd | Negative | Negative |
| | | | Serum | ZD/25/7-Se | Negative | Negative |
| | | | Nasal Swab | ZD/25/7-NS | Negative | Negative |

**Appendix Table 2. Identification of the closest hits to HPAI A(H5N1) clade 2.3.2.1a viruses isolated from two cats (A/Cat/Chhindwara/ZD25-1/2025 & A/Cat/Chhindwara/ZD25-2/2025) through nucleotide BLAST analysis against the complete GISAID database.**

| Gene segments | Closest hit on nucleotide blasting with entire GISAID database | Identity | Clade classification |
|---|---|---|---|
| PB2 | A/Victoria/149/2024 (A/H5N1) | 99% | 2.3.4.4b |
| PB1 | A/Victoria/149/2024 (A/H5N1) | 98% | 2.3.4.4b |
| PA | A/Victoria/149/2024 (A/H5N1) | 99% | 2.3.4.4b |
| HA | A/Victoria/149/2024 (A/H5N1) | 99% | 2.3.2.1a |
| NP | A/Victoria/149/2024 (A/H5N1) | 98% | 2.3.2.1a |
| NA | A/Victoria/149/2024 (A/H5N1) | 99% | 2.3.2.1a |
| MP | A/Bean goose/Korea/21WC198/2022 (A/H5N1) | 99% | 2.3.4.4b |
| NS | A/Victoria/149/2024 (A/H5N1) | 99% | 2.3.2.1a |

**Appendix Table 3. Amino acid substitutions identified in different proteins encoded by the HPAI A(H5N1) clade 2.3.2.1a viruses isolated from two cats in comparison to A/Victoria/149/2024 (A/H5N1)**

| **Genes** | **AA position** | **A/Victoria/149/2024** | **A/Cat/Chhindwara/ZD25-1/2025** | **A/Cat/Chhindwara/ZD25-1/2025** |
|---|---|---|---|---|
| HA | 145 (129-H5 numbering) | L | R | R |
| | 242 (226-H5 numbering) | I | V | V |
| | 402 (387-H5 numbering) | D | N | N |
| | 519 (504-H5 numbering) | S | N | N |
| PB2 | 590 | G | S | S |
| PB1 | 97 | G | E | E |
| | 121 | R | K | K |
| | 211 | R | K | K |
| | 383 | **E** | **D** | **E** |
| | 525 | I | V | V |
| | 581 | G | E | E |
| | 688 | M | V | V |
| PB1-F2 | 66 | **G** | **S** | **S** |
| | 90 | D | N | N |
| PA | 262 | K | R | R |
| | 327 | K | E | E |
| | 479 | E | D | D |
| | 492 | K | R | R |
| | 505 | I | V | V |
| | 607 | M | V | V |
| NP | 400 | R | K | K |
| | 475 | V | I | I |
| NA | 34 | V | I | I |
| | 42 | L | F | F |
| | 181 | G | S | S |
| | 221 | I | V | V |
| NS | 74 | D | N | N |

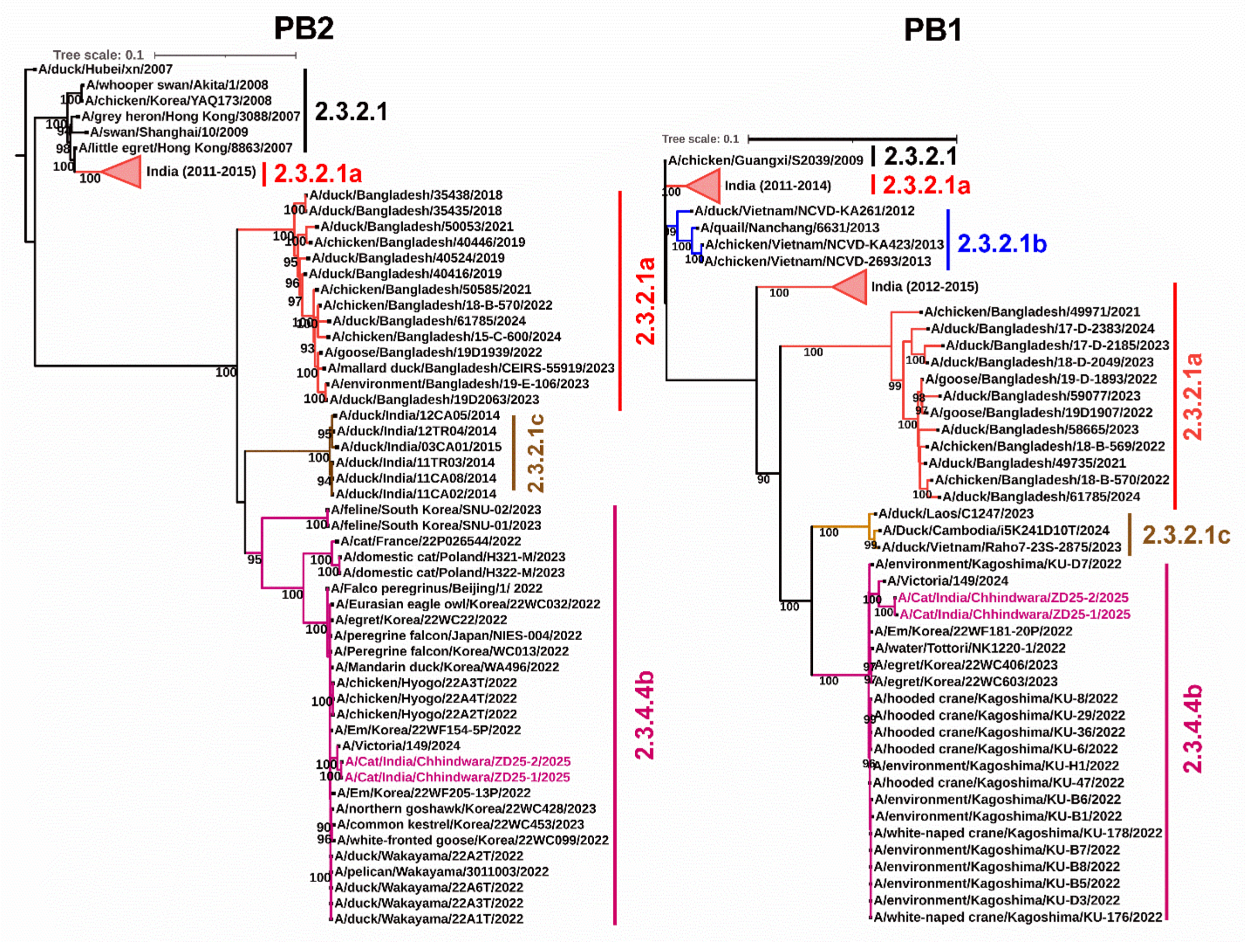


**Appendix Figure 1.** ML tree showing the phylogenetic relationship among the cats HPAI H5N1 PB2 and PB1 sequences of this study with closely matched H5N1 clade 2.3.4.4b and different representative clade (2.3.2.1, 2.3.2.1a, 2.3.2.1b, and 2.3.2.1c) sequences from the GISAID database.

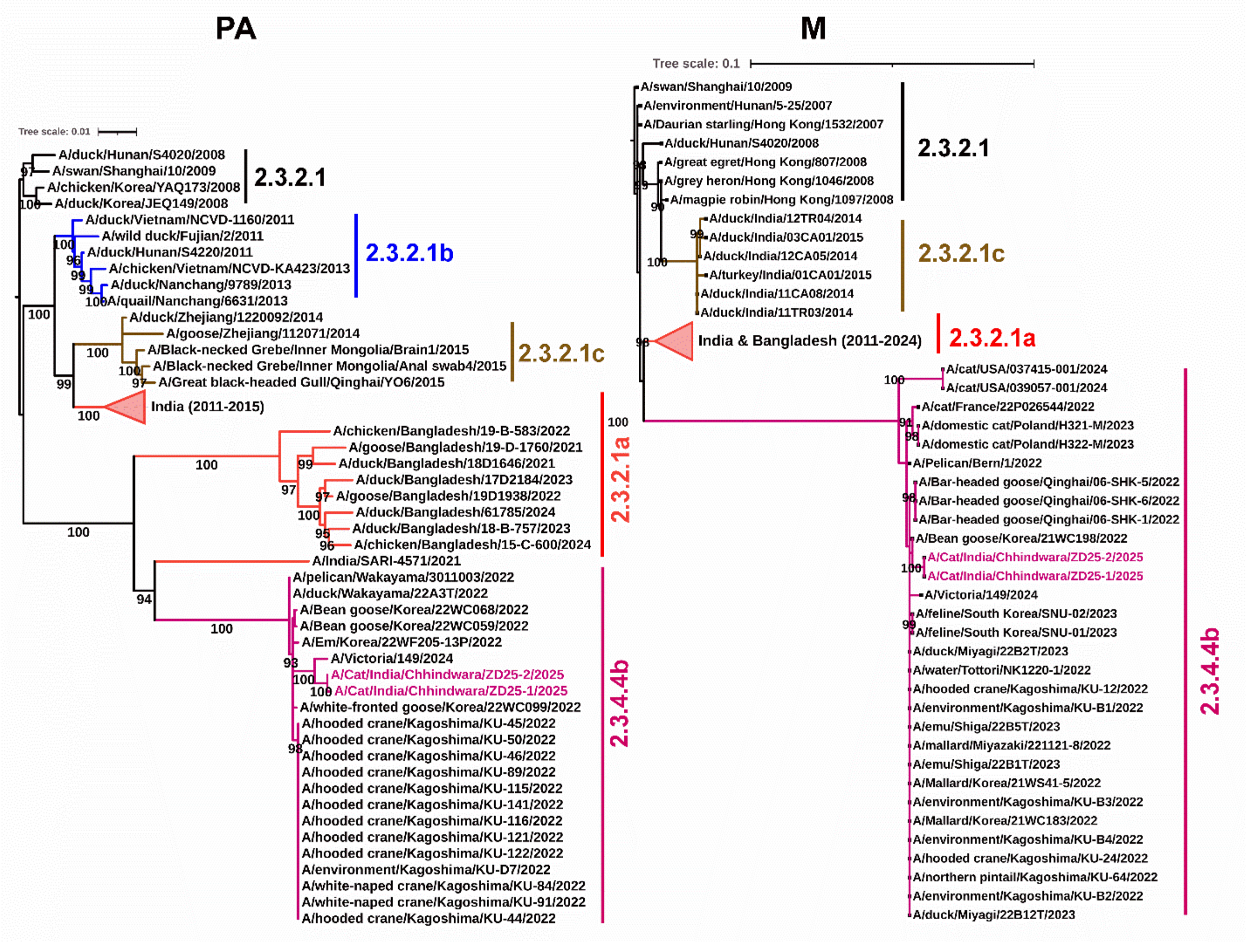


**Appendix Figure 2.** ML tree showing the phylogenetic relationship among the cats HPAI H5N1 PA and MP sequences of this study with closely matched H5N1 clade 2.3.4.4b and different representative clade (2.3.2.1, 2.3.2.1a, 2.3.2.1b, and 2.3.2.1c) sequences from the GISAID database.

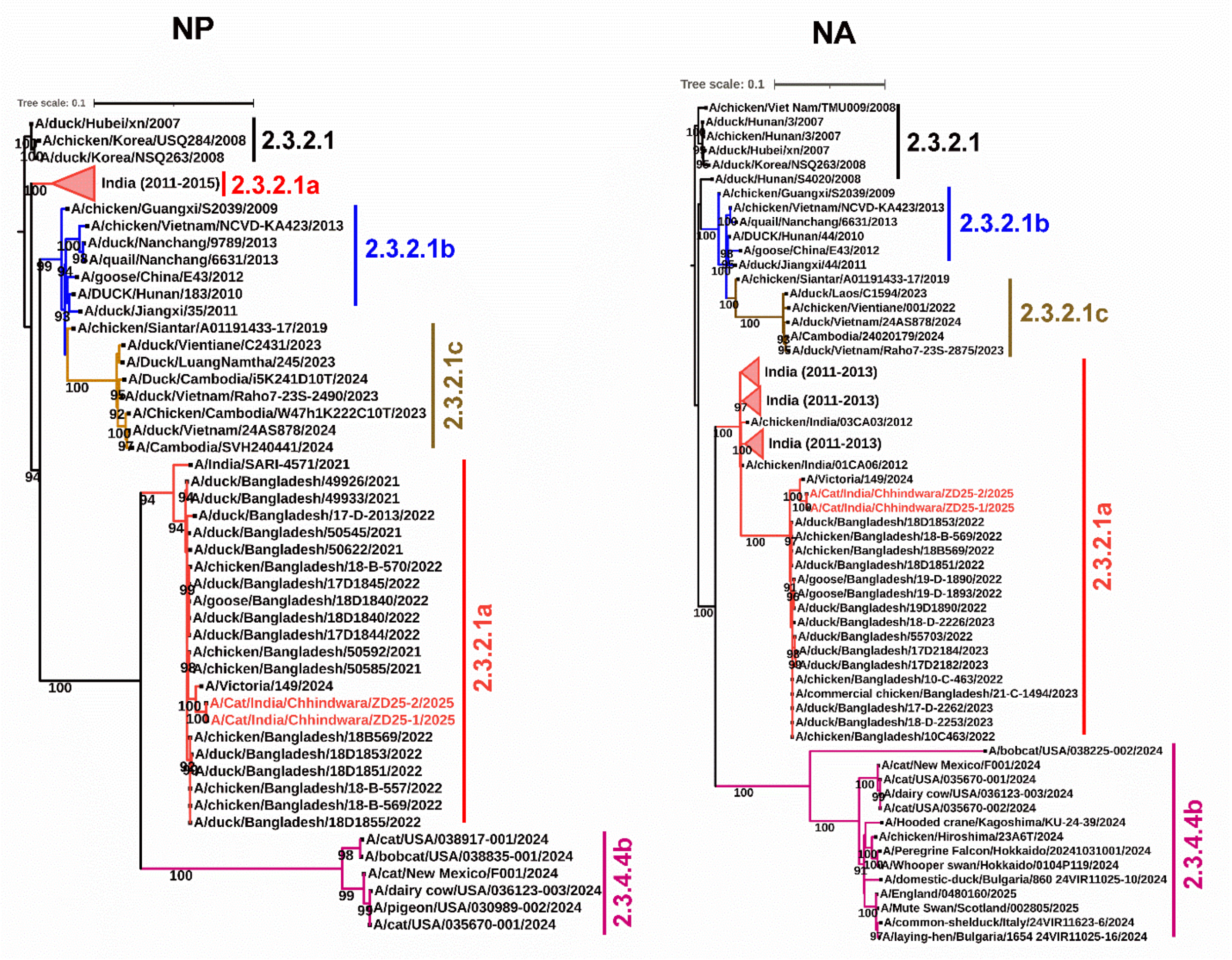


**Appendix Figure 3.** ML tree showing the phylogenetic relationship among the cats HPAI H5N1 NP and NA sequences of this study with closely matched H5N1 clade 2.3.2.1a and different representative clade (2.3.2.1, 2.3.2.1b, 2.3.2.1c, and 2.3.4.4b) sequences from the GISAID database.

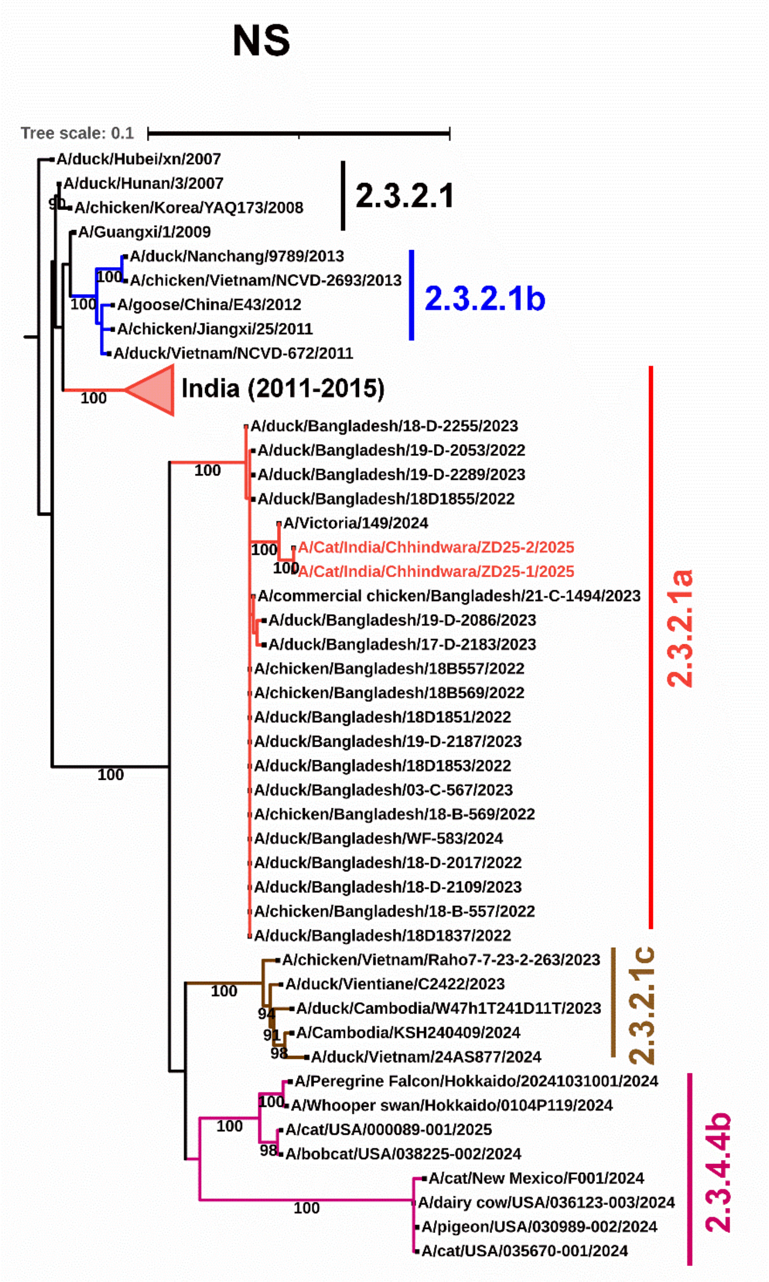


**Appendix Figure 4.** ML tree showing the phylogenetic relationship among the cats HPAI H5N1 NS sequences of this study with closely matched H5N1 clade 2.3.2.1a and different representative clade (2.3.2.1, 2.3.2.1b, 2.3.2.1c, and 2.3.4.4b) sequences from the GISAID database.
